# scoup: Simulate Codon Sequences with Darwinian Selection Incorporated as an Ornstein-Uhlenbeck Process

**DOI:** 10.1101/2025.06.14.659628

**Authors:** Hassan Sadiq, Darren P. Martin

## Abstract

Genetic analyses of natural selection within and between populations have increasingly developed along separate paths. The two important genres of evolutionary biology (i.e. phylogenetics and population genetics) borne from the split can only benefit from research that seeks to bridge the gap. Simulation algorithms that combine fundamental concepts from both genres are important to achieve such unifying objective. We introduce scoup, a codon sequence simulator that is implemented in R and hosted on the Bioconductor platform. There is hardly any other simulator dedicated to genetic sequence generation for natural selection analyses on the platform. Concepts from the Halpern-Bruno mutation-selection model and the Ornstein-Uhlenbeck (OU) evolutionary algorithm were creatively fused such that the end-product is a novelty with respect to computational genetic simulation. Users are able to seamlessly adjust the model parameters to mimic complex evolutionary procedures that may have been otherwise infeasible. For example, it is possible to explicitly interrogate the concepts of static and changing fitness landscapes with regards to Darwinian natural selection in the context of codon sequences from multiple populations.

## Statement of need

Statistical inference models that are used to analyse the degree of the impact of Darwinian natural selection on observed genetic data, command a healthy portion of the phylogenetic literature (Gupta and Vadde 2023). Validation of these largely codon-based models relies heavily on simulated data. Given the ever increasing diversity of natural selection inference models that exist (Yang 2007; Arenas 2015; Kosakovsky Pond et al. 2020), there is a need for more sophisticated simulators to match the expanding model complexities.

Bioconductor (Gentleman et al. 2004) is a leading bioinformatics platform distributing peer-reviewed R packages. A search of the entries on the platform, in Version 3.22 on 18 February 2026, with keywords including, `codon, mutation, selection, simulate`, and `simulation` returned a total of 70 packages (excluding scoup) out of the 2361 available. None of the retrieved entries was dedicated to codon data simulation for natural selection analyses. Thus, `scoup` is designed on the basis of the mutation-selection (MutSel) framework (Halpern and Bruno 1998) as an overdue contribution to the void.

Software and/or packages for simulating molecular protein sequences are a few in the scientific literature (Peng et al. 2015). Existing simulators tend to be more suitable for quantitative character evolution. These include, `ape` (Paradis and Schliep 2019), `ouch` (Cressler, Butler, and King 2015; Butler and King 2004) and `geiger` (Pennell et al. 2014). Other extensively used DNA sequence simulators including, `Seq-Gen` (Rambaut and Grass 1997), `INDELible` (Fletcher and Yang 2009), `PhyloSim` (Sipos et al. 2011) and `phangorn` (Schliep 2011) are parameterized in accordance with *ω*-based models (Goldman and Yang 1994; Muse and Gaut 1994). More recent sequence simulators, such as, `phastSim` (Maio et al. 2022) and `AliSim-HPC` (Ly-Trong, Barca, and Minh 2023) prioritized output capacity. Only few genetic simulators were built upon the more elaborate MutSel evolutionary concept. These include, `Pyvolve` (Spielman and Wilke 2015a) and `SGWE` (Arenas and Posada 2014). To the best of our knowledge, these existing MutSel friendly simulators are only able to generate data from static landscapes. With our proposed simulator, it is possible to generate codon sequences from landscapes that are static or those that are changing (also known as *seascapes*) (Mustonen and Lässig 2010).

## Algorithm

`scoup` is further unique for at least three reasons. First, it incorporates Darwinian natural selection into the MutSel model in terms of variability of selection coefficients, an extension of an idea from Spielman and Wilke (2015b). Second, it directly utilises the concept of fitness landscapes. Third, fitness landscape updates can be executed in either a deterministic or a stochastic format. The stochastic updates are implemented in terms of the more biologically amenable, Ornstein-Uhlenbeck (OU) process (Bartoszek et al. 2017; Uhlenbeck and Ornstein 1930). A crude summary of how substitution events are executed in scoup is presented in Figure 1.

**Figure 1:**
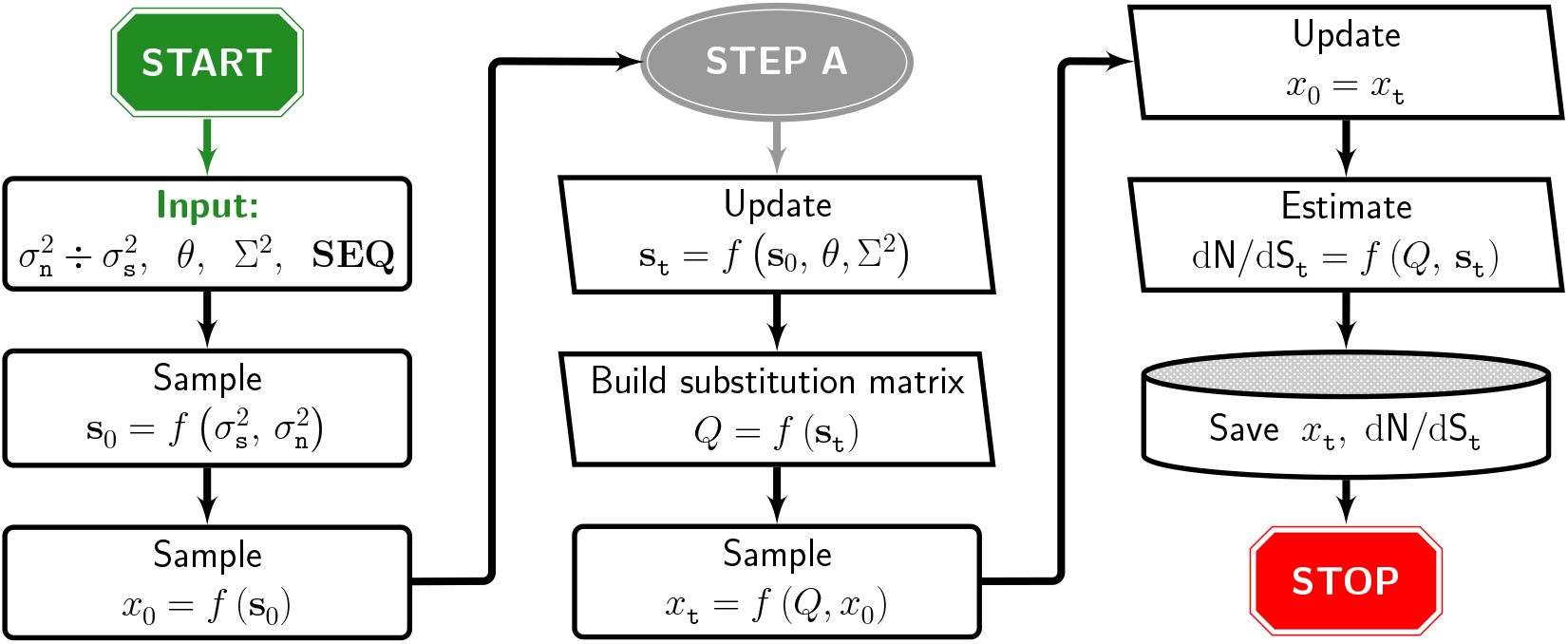
Summarised scoup algorithm. The flowchart shows the process for a single substitution event. After each substitution event, the process returns to *STEP A*, until the input tree length (*τ* ∈ **SEQ**) is exhausted. 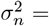 variance of amino acid selection coefficients. 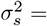 variance of synonymous codon selection coefficients. Σ^2^ = OU asymptotic variance. *θ* = OU mean reversion rate. **SEQ** = sequence information. *x*_⋆_ = codon. **s**_⋆_ = codon selection coefficient vector.

We highlight two important design choices from Figure 1. First, we assume that a static fitness landscape is obtained from a single set of parameters (*ξ*) needed to sample a 20-element numerical vector of amino acid selection coefficients (that is, *s*_0_ in Figure 1). The coefficients are subsequently used as inputs of the corresponding MutSel model. The seascape setting is then defined as a function of multiple sets of parameters (*ξ*_1_, *ξ*_2_, …, *ξ*_*k*_, for *k* ≤ extant taxa size). Second, the coefficient update (*s*_*t*_) step is done after every substitution event. In addition, the Ornstein-Uhlenbeck update process is discretised. In other words, the OU jump sizes are fixed and pre-specified as an input to the simulation functions.

## Implementation

`scoup` is primarily designed using base functions in R. Some important complementary functions are imported from the `Matrix` (Bates, Maechler, and Jagan 2024) and the `Biostrings` (Pagès et al. 2024) packages. We simulated some sequences with scoup to verify the accuracy of the outputs from the package. The output data each comprise eight sequences and 1000 codon sites. All the other necessary model parameters were kept the same for all simulated replicates. The data and all the associated files, including the simulation and analyses code, are available in the `paper/data` folder as part of the package. Only the variance of the selection coefficients of the synonymous codons 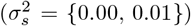 and the variance of the amino acids 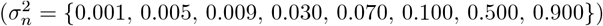 were varied and five replicate sequences were generated for each 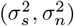 combination. The data sets were analysed with PAML (Yang 2007) to obtain maximum likelihood estimates of the ratio of the rates of non-synonymous to synonymous substitutions (*ω*) and these were compared to the analytical estimates (d*N/*d*S*) obtained from `scoup`. The results are summarised in Figure 2.

**Figure 2:**
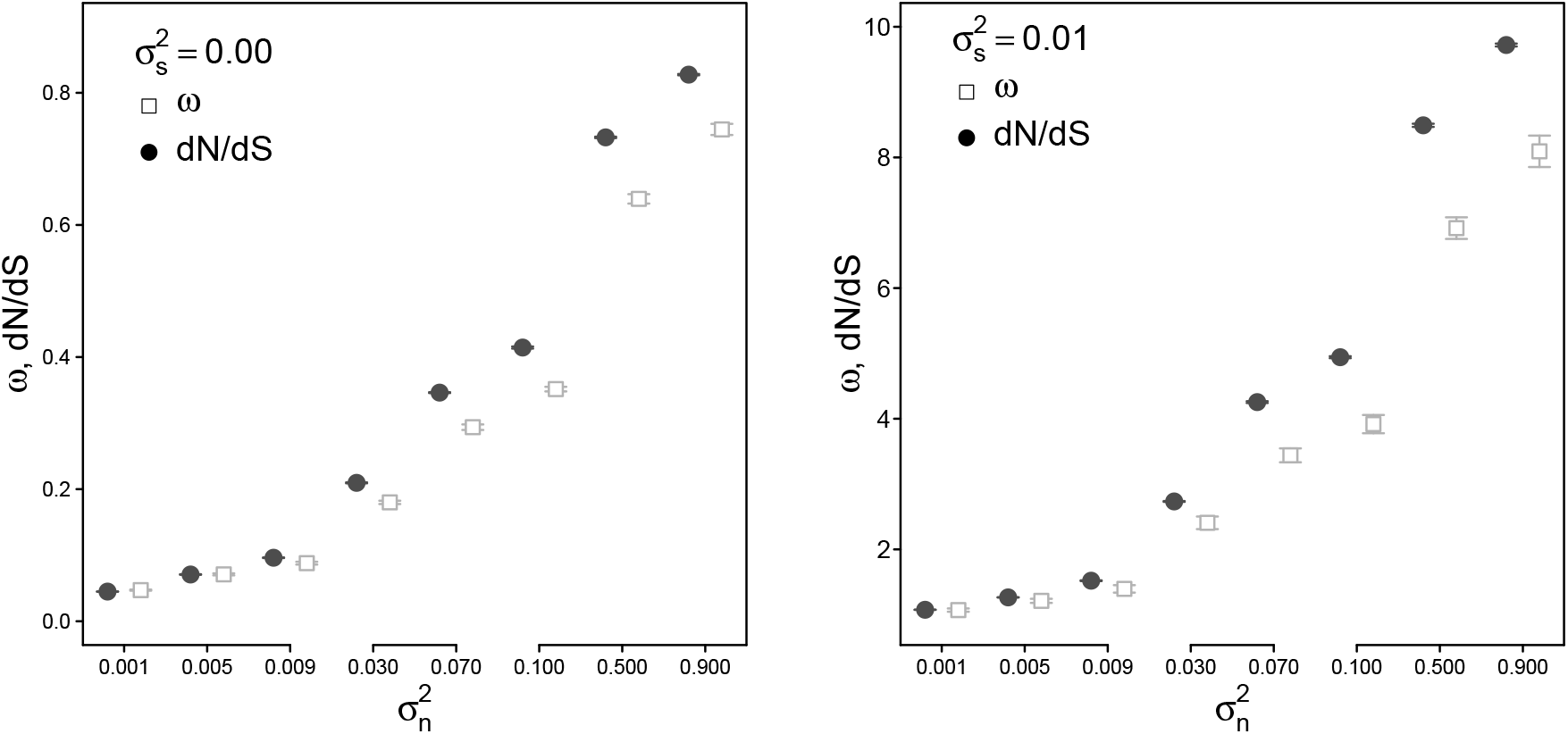
Analyses of data generated with scoup. The width of the arrows correspond to two times the associated standard errors. 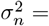 variance of amino acid selection coefficients. 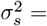 variance of synonymous codon selection coefficients. *ω* = maximum likelihood estimate of non-synonymous to synonymous substitution rates ratio obtained using PAML, d*N/*d*S* = analytical equivalent of *ω* that is returned as part of the outputs from `scoup`.

Three features from Figure 2 are noteworthy. First, there is good correlation between the simulated (as measured by d*N/*d*S*) and the inferred (as measured by *ω*) magnitude of natural selection effect. The slight discrepancies as 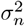 increases are likely due to the limited sizes of the data sets (Spielman and Wilke 2015a). This imply that outputs from scoup are reliable. Second, as expected (Spielman and Wilke 2015b), in the absence of synonymous selection (that is, 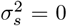) the selection effect is predominantly negative (that is, d*N/*d*S, ω <* 1) and the effect is largely positive when synonymous selection is present. This further asserts of the reliability of the outputs from `scoup`. Third, the magnitude of natural selection effect may be influenced by amino acid selection (or aptly, non-synonymous selection). This property is yet to be thoroughly investigated in the computational molecular evolution literature and there is hardly any other available computational resource that permits its exploration. This underlines the potential importance of `scoup`.

## Conclusions

We present scoup, a R package for codon sequences simulation, where the evolutionary processes are mirrored more realistically than most existing simulators. Our framework creatively incorporates the Ornstein-Uhlenbeck process into the mutation-selection evolutionary model. This attribute could potentially unlock exciting research avenues that will improve existing knowledge about the complex interactions of different, potentially interacting, molecular evolutionary processes.

## Code availability

`scoup` is published for free public use under the GPL-2 license. It is available for download from the Bioconductor platform, along with detailed documentation and tutorial files. Some additional sample code are accessible in the `tests` and the `vignettes` folders of the package.

## Acknowledgements

We thank Ben Murrell for suggesting modelling varying selection coefficients with an OU process. Computations were performed using the `HPC1` facility at Stellenbosch University, South Africa.

